# Measuring developmental information encoded by a dynamical landscape

**DOI:** 10.64898/2026.03.03.709461

**Authors:** Meritxell Sáez, Giorgos Minas, Elena Camacho-Aguilar, David A. Rand

**Affiliations:** Department of mathematics and data analysis, IQS, Universitat Ramon Llull, Via Augusta 390, 08017 Barcelona, Spain; School of Mathematics and Statistics, University of St Andrews, St Andrews, KY16 9SS, Scotland, United Kingdom; Department of Gene Regulation and Morphogenesis, Centro Andaluz de Biología del Desarrollo (CSIC-UPO-JA), 41013 Seville, Spain; Mathematics Institute & Zeeman Institute for Systems Biology and Infectious Epidemiology Research, University of Warwick, Coventry CV4 7AL, United Kingdom

**Author notes:** These authors contributed equally to this work (Meritxell Sáez, Giorgos Minas).

**Keywords:** Information theory, Landscape models, Decision-making systems

## Abstract

During embryogenesis, as cells proliferate and assemble into tissues, they undergo a sequence of transitions between distinct molecular states eventually giving rise to a cellular population consisting of an appropriate distribution of specific functional cell types. Recent progress on the dynamics underlying decision-making in developmental landscape makes it feasible to start analysing the amount of information involved in constructing such systems. To explore this we introduce the notion of potency of a developmental landscape and attempt to calculate it for two development systems of current interest, in-vitro differentiation of epiblast-like cells into neural and mesodermal progenitors and the worm vulva patterning system. Our approach integrates concepts from developmental biology, information theory and dynamical systems to estimate both the number and identity of signalling regimes that give rise to distinguishable temporal response patterns.

---

The transmission and processing of information is crucial for the function of all living systems [19], and information theory [8] is currently the only framework for defining, quantifying and rigorously analysing this otherwise intractable notion. While information theory has been successfully used to study intracellular signaling at the molecular level [20, 24, 17, 2, 5, 18] and to the genetic coding of positional information of cells in developing embryos [9, 22, 13, 21], much less work has focused on applying these concepts to other levels of systems organisation. These include processes such as the allocation of cells to distinct cell states and fates during embryogenesis, or the organisation of these states into functional patterns. In this paper we consider the process— often termed cellular decision making—whereby, during embryonic development, multipotent progenitor cells generate the appropriate and diverse distribution of specialised, functional cell types. To study this process we use information theory to measure the ability (which we call *potency*) of this decision making machinery, or parts of it, to produce recognisably distinct outcomes.

Recent work has formalised an approach to describe such processes using dynamical systems theory, which opens up the opportunity to analyse single cell data from such systems and quantify information in this context. In this approach [7, 3, 15, 16, 14, 6] cell states correspond to attractors in a dynamical landscape and fate decisions emerge from signal-induced changes to the landscape topology. Cells transition between states following paths defined by saddle points while being subject to the intrinsic noise underlying the system. This formulation provides a mathematical link between the stochastic dynamics defined by the underlying gene regulatory networks (GRNs) and cellular behaviour in terms of cell states and transitions between these states. The information carried by this process can therefore be characterised by how effectively the GRN, modulated by signals, can generate distinct cell outcomes.

It has been shown in [7, 3, 15, 10, 6] that experimental single cell data can be used to construct and estimate the parameters of landscape models that can reproduce the data quantitatively. The availability of these data-calibrated generative models allows us to develop a method for calculating information quantities associated with these developmental systems.

Here, we present a practical and interpretable method to estimate the informational quantities described below for two exemplar developmental systems: the in-vitro differentiation system of Epiblast-like cells into neural and mesodermal progenitors [15] and the spatial patterning of the worm vulva [3]. By using the calibrated landscape models of these systems we produce a large number of time-trajectories for these systems, reflecting the underlying GRN dynamics and intrinsic noise. The high dimensionality, and the correlations in our variable data pose a challenge for the estimation problem. By adopting a pragmatic approach in summarising the data and tailoring to our setting ideas from [4] we are able to estimate the quantities of interest for both systems.

By estimating these quantities, our framework provides a way to ask how many —and which— signaling regimes are expected to produce recognizably distinct outcomes, and which are effectively redundant. This analysis can guide the design of experiments that seek to identify signals or combinations of signals capable of modulating cell outcomes, as well as experiments aiming to uncover redundancy or robustness in developmental systems. In particular, we show how our notion of information transfer can identify regimes that cells can, or cannot, reliably distinguish, and quantify the confusion introduced when indistinguishable regimes are added to the system. Furthermore, these predictions yield quantitative estimates of information transfer that can be tested against future data and used to guide model refinement.

After introducing the concept of potency and simple motivating examples in Section 1, we present a minimal landscape model to illustrate the approach in 2 and describe our estimation procedure in Section 3. We then apply the method to the in-vitro differentiation system of Epiblast-like cells into neural and mesodermal progenitors and to the spatial patterning of the worm vulva in Sections 4 and 5. The former example deals with decision making driven by different morphogen combinations, while the latter looks at experimental perturbations affecting signaling and the resulting cell fate patterning. These examples illustrate how the approach can be applied in different developmental contexts.

## 1 Potency and mutual information

In our examples, for each sampling of the decision-making process we obtain a temporal record *R* of the distribution of cell states in the population at multiple time-points. We call this the *temporal response pattern* (TRP). In general, this would be characterised by some set of summary statistics. These might, for example, contain the proportion of cells in each cell state at different times and also perhaps information on their position or spatial distribution in different parts of the embryo. We aim to calculate the number of distinguishable TRPs generated by a given set of signaling regimes *𝒮* and will call this quantity the *potency*. A simple example inspired by our data is shown in Fig. 1 where just four signals are considered and a 2-dimensional projection of the responses *R* is shown. In the main examples these responses will be high dimensional vectors giving summary statistics of the population at several time points. Looking at the right panel we see that two of the responses are clearly distinguishable from the others but the other two are hardly distinguishable from each other. If one of the signals, called *S*_3_ or *S*_4_, were not considered, the three remaining signals would be clearly distinguishable and the potency would be 3. With all four signals present, it is less clear what the potency should be, but the method we present below allows us to approximate it.

**Figure 1:**
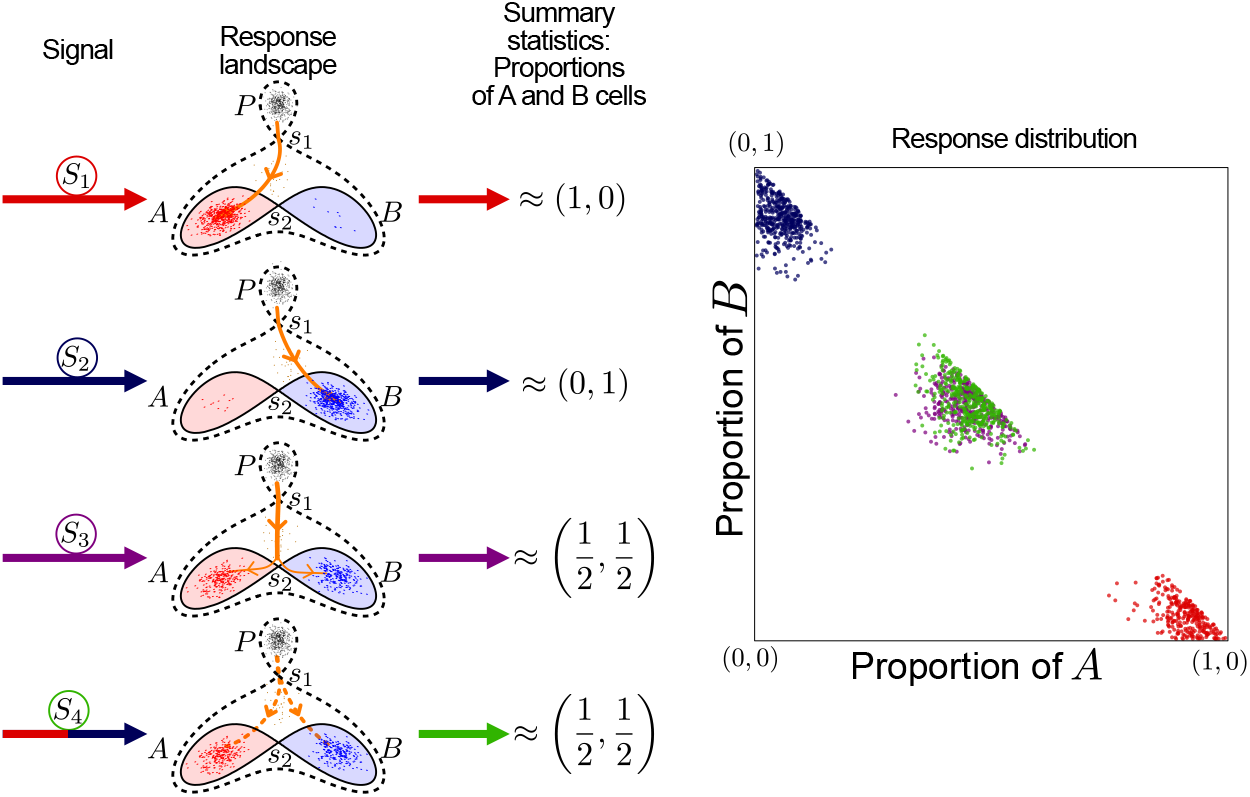
Binary flip model for decision-making. Cells begin in a progenitor state (black) and can receive signals *S*_1_ or *S*_2_, leading to type *B* cells (blue) or type *A* cells (red). If they receive signal *S*_3_, they become type *A* or *B* with equal probability (1/2). A timed combination of signals *S*_1_ and *S*_2_ (labelled *S*_4_) give a similar response to *S*_3_. This decision-making process can be modelled by the *binary flip landscape* of [15], which features three attractors and two saddles. Various signals bifurcate the top attractor (progenitor state, P) and move the unstable manifold of the top saddle, prompting cells to transition from *P* to one of the bottom attractors *A* or *B*. The response to this system can be summarised by the proportion of cells of types A and B at a certain time point. Repeating the process several times and plotting the responses in the proportion space (right panel) one can see three clusters of responses with signals 1 and 2 clearly separated and signals 3 and 4 laying one on top of the other. Note that the presence of cells still in transition at response time causes the proportions to sum to less than one in many cases.

Generally, this TRP *R* is stochastic and will vary from one repetition of the cell decision-making process to another. Therefore, to the input signal *S*, we associate a probability distribution *P*_*S*_(*R*) from which the TRP *R* is sampled. This relation *S* →*P*_*S*_(*R*) between the signal and the probability distribution of the response defines what in information theory would be called a *channel*. In addition to the channel, a distribution *P* (*S*) over the signals may be specified, defining the probability with which each signal is used.

We regard the responses *R* to the signals *S* ∈ *𝒮* as being distinguishable if the distributions *P*_*S*_(*R*), S ∈ *𝒮*, are not packed too tightly so that by monitoring the response *R* we can reliably infer which signal *S* was used. Mathematically, we interpret this loose packing as corresponding to the property that for all pairs of signals *S, S*^*′*^ in *𝒮*, when *R* is drawn from *P*_*S*_, 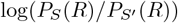 is large. This corresponds to demanding that the relative entropy or Kullback-Leibler divergence 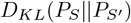 of *P*_*S*_ and 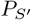 is large where, in general, *D*_*KL*_(*P* ||*Q*) is the *P*-mean of log_2_ *P* (*x*)*/Q*(*x*). Rather than using 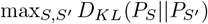 to measure how spread out these distributions are, it is better to use the infimum of the quantity 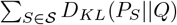 over all distributions *Q* [26, 12] which is small when the probabilities are tightly packed and large when they are spread out. A way to see this is to note that if *P*_*S*_ is disjoint from the other response distributions and we take *Q* to be the equal weight mixture of all response distributions, then *D*_*KL*_(*P*_*S*_ ||*Q*) is log_2_ |*𝒮*|, which is the maximum possible value.

If we now consider a finite set of *N* signals, *𝒮*, this quantity is minimised by taking for *Q* the equal weight mixture 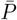 of the *P*_*S*_, *S* ∈ *𝒮* [26, 12]. This is proportional to the well-known Jensen-Shannon divergence

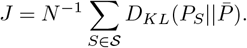

This is equal to the mutual information (MI), *I*(*S*; *R*), between *S* and *R* for the channel where all the signals are equally likely. It has some interesting properties, for example,

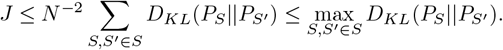

Moreover, it is directly related to how many of the responses we can distinguish by the following result [26]: for any finite set {*Q*_*α*_ : *α* ∈ *G*} of probability measures,

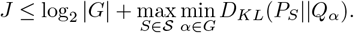

The |*G*| probability measures *Q*_*α*_, *α* ∈ *G* can be viewed as an approximation of the *N* probability measures *P*_*S*_, *S* ∈ *𝒮*. The term max_*S*_ min_*α*_ *D*_*KL*_(*P*_*S*_ || *Q*_*α*_) then denotes the approximation error, measured via the Kullback-Leibler divergence. The right hand side of this inequality can therefore be made small if it is possible to choose not too many probability measures *Q*_*α*_ which well approximate the given set of probability measures *P*_*S*_. This motivates our definition of *potency* as

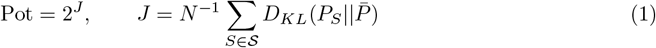

We can also consider this from a minimum risk point of view and ask about obtaining lower bounds for the minimum risk when using estimators *T* such as machine learning classifiers. When the TRPs corresponding to the different signals are closely packed it is hard to distinguish between them and hence, the risk of misclassification will be large. The minimum risk *r* = inf_*T*_ sup_*S*∈*𝒮*_ *P*_*S*_(*T* ≠ *S*) which is bounded below by

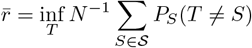

and then 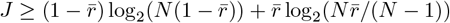. Thus we have (see [12] for more details)

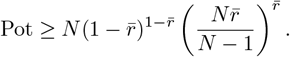

Given this approach, the potency of the example in Fig. 1 is easy to calculate as the distributions 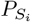 are proportional to 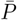 when restricted to the support of 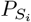. Therefore, 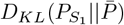 and 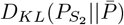 are both equal to log_2_ 4 = 2 while 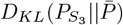) and 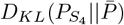 are both equal to log_2_ 4*/*2 = 1. Thus the potency is 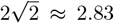. If we consider only the first 3 signals, then 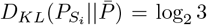 for *i* = 1, 2, 3 and hence the potency is clearly 3. The former is clearly well below the maximum potency 4 and even below 3 which would apply if either the signal *S*_3_ or *S*_4_ were left out. This reflects that we can easily decide whether one of *S*_3_ or *S*_4_ took place but not which.

If the signals *S* ∈ *𝒮* are sampled according to weights *w*_*S*_ (*w*_*S*_ *>* 0.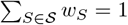) then the above discussion applies with 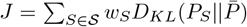, which is again equal to the MI, *I*(*S*; *R*), now for the channel with weighed signals. We limit ourselves to three different possibilities for such weights which correspond to *in vivo*, experimental and maximal potency situations. In the first, the weights are determined by the processes taking place *in vivo*. The second is where an experimenter chooses how to apply the signals, and, for this case, we will assume that all the signals are applied with equal probability. In the third case we would like to determine the distribution of signals which maximizes potency. Currently it is not feasible to estimate the weights in the *in vivo* case but the potency for the optimal case gives an upper bound for this.

## 2 Confusion in landscape models for cell decision making

Consider a simple decision-making system where cells begin in an initial progenitor state and respond to signals that direct their fate. Signal *S*_1_ (e.g., low morphogen concentration) leads to type A cells, while *S*_2_ (e.g., high concentration) results in type B cells. Signal *S*_3_ (e.g., medium concentration) causes cells to become either type A or B randomly with equal probability.

This decision-making system can be modeled using a binary flip landscape (Fig. 1), which includes three attractors—*P* (progenitor), *A*, and *B*—and two saddles, *s*_1_ and *s*_2_. Cells begin at the top attractor and follow stochastic dynamics defined by the underlying landscape. Signal *S*_1_ (or *S*_2_) destroys the progenitor state via collision with saddle *s*_1_, causing cells to move along an orange escape path to attractor *A* (or *B*). With signal *S*_3_, the path passes near saddle *s*_2_, and noise causes cells to reach either *A* or *B* with equal probability. In all cases, the escape path follows the unstable manifold of *s*_1_, with noise introducing variability in cells trajectories.

Defining the system’s response as the vector of proportions of type *A* and *B* cells at a given time, the responses to signals *S*_1_, *S*_2_, and *S*_3_ are approximately (1, 0), (0, 1), and 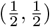, respectively. Due to noise and incomplete transitions, these values may vary and not sum to 1. For simplicity, a cell is considered to have reached an attractor once it enters the shaded region. Repeating the process for several cell populations and plotting the results yields three distinct response clusters as in Fig. 1 (right panel), giving the system a potency of 3.

We consider now a fourth signaling regime consisting in applying signal *S*_1_ until 50% of the cells are *A* and then switching to signal *S*_2_ for the remaining of the time course. The response for this regime will be again roughly 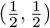. When we look at several repetitions of the process, the vectors obtained overlap those corresponding to *S*_3_, hence it is impossible to distinguish, given a response close to 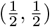 which signaling regime produced it. The potency of the system for these four signals is 2.83. Thus in adding a signal we have decreased potency. This we call *confusion*, and it is discussed below.

It is important to note that the responses to signalling regimes *S*_3_ and *S*_4_ may be distinguishable at time points other than the one considered above. For this reason, our approach uses the TRP to compute potency, trying to ensure that any observed confusion is not an artifact of the chosen observation times.

We also emphasize that we are not tracking here the fate of individual cells, and therefore we do not care which specific cell ends up in which cluster. If we were to track individual cells, and there were *n* cells, then the analogue of the potency would be of the order 2^*nJ*^ rather than 2^*J*^ because we would be specifying the fate of each cell.

## 3 Estimating potency

Calculating the potency will not be easy in general as the responses for developmental systems will typically be high dimensional and highly correlated. Since direct methods for calculating MI are not well suited for this type of data, we take a different approach, introduced in [4], that can give a good estimate of the lower bound of the MI for our systems. This approach uses machine learning methods, specifically supervised classification, for the estimation of the information transmitted. The method treats the different signals in *𝒮* = {*S*_1_, *…, S*_*N*_} as labels and trains a set of classifiers *χ* to predict the label from a given response *R*. A classifier is a map from the space of responses to the space of signals, i.e. *Ŝ* = *χ*(*R*). As a function of observations that are randomly sampled, *Ŝ* = *χ*(*R*) is also a random variable and has an associated probability distribution. In our systems *S* and hence *Ŝ* are discrete variables and *R* is continuous and high dimensional so it is much easier to calculate *I*(*S*; *Ŝ*) than *I*(*S*; *R*). Moreover, *I*(*S*; *Ŝ*) is a lower bound for *I*(*S*; *R*). Indeed, the chain rule for mutual information implies that

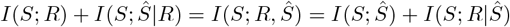

where *I*(*S*; *Ŝ*|*R*) is the (remaining) mutual information between *S* and *Ŝ* given a value of *R* [8]. But *I*(*S*; *Ŝ*|*R*) = 0 since *R* completely determines *Ŝ* via the classifier, and therefore

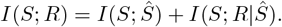

The development of computationally efficient and accurate classifiers provides the potential that *I*(*S*; *R*|*Ŝ*) can be made small so that the method can provide estimates close to the actual MI, particularly when we have large amounts of observations. In this case, we get that 2^*I*(*S*; *Ŝ*)^ ≈ 2^*I*(*S*;*R*)^ = Pot.

We will need to sample a *training data set* from *P* (*S, R*) of fixed size to parameterise various classifiers and indicate which of them performs best. The best performing classifier will depend on the specific structure of the data considered (see below and SI). Given such a classifier, we will estimate *I* = *I*(*S*; *Ŝ*) by sampling a *test data set* disjoint from training. We will perform this estimation many times, sampling new data sets (both training and test), to obtain an empirical distribution *P* (*I*) for *I*.

This gives not only the median as an estimate for *I* and the potency 2^*I*^ but also a confidence interval for it which can be shrunken by increasing the number of repetitions, *m*, of this process. It is quite possible that the performance of the training data set is in the right-hand tail of *P* (*I*) so that using it to estimate *I* would lead to overestimation. Therefore, for an effective estimate with a small confidence limit, in addition to the need for extensive training data, one needs a large number of test data samples different from the training ones: that is, a large number of repeats of the process by which a given signal generates a given TRP (see SI, section 1 for details). These would be very costly to obtain only from experimental data. This is where the availability of a dynamical landscape model is key since it allows the generation of as many replicates of the experiment as needed while reproducing the variability of the experimental setting (Fig. 2D).

**Figure 2:**
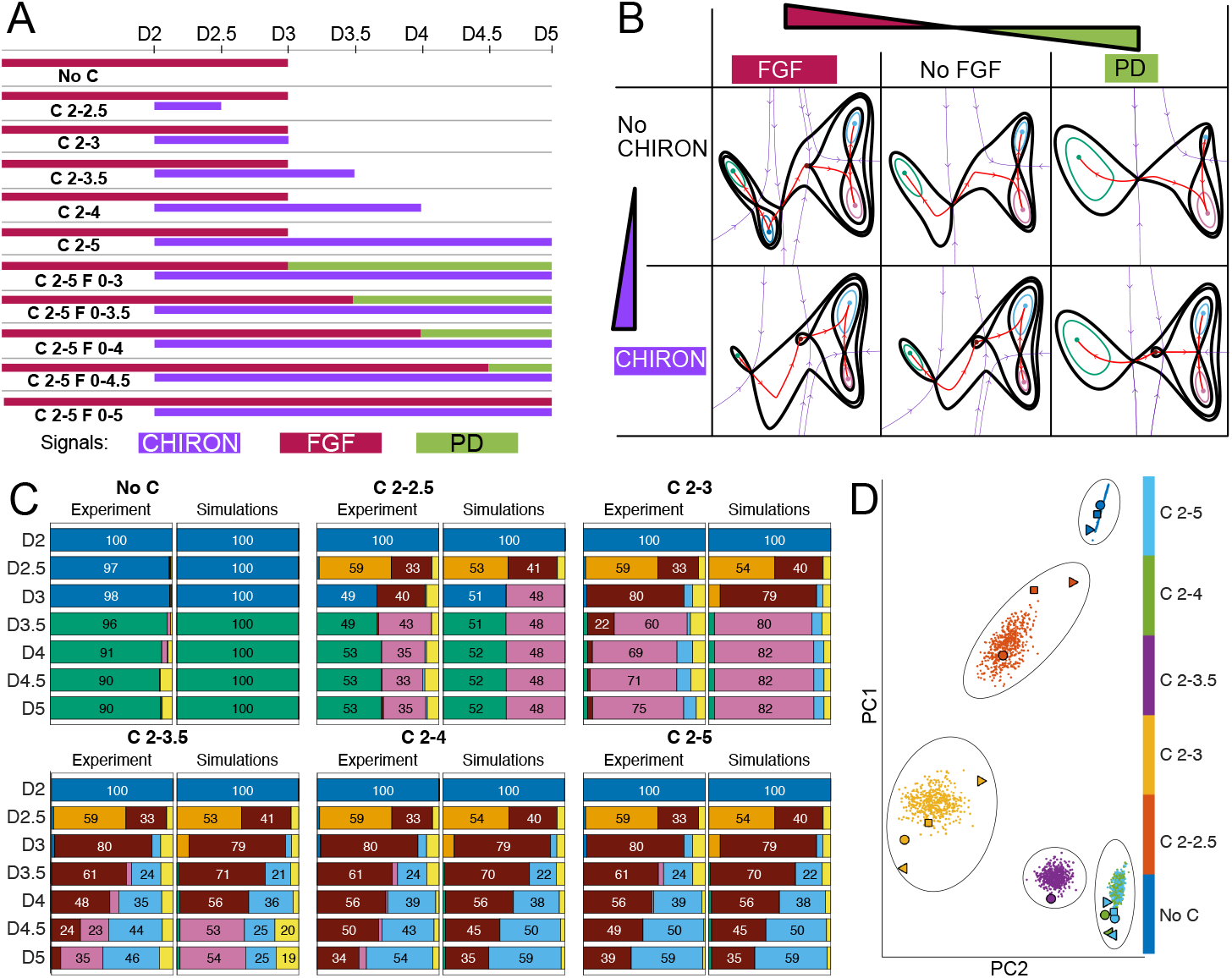
*Adapted from [15]*. A model for the in-vitro differentiation of ES cells into neural and mesodermal derivatives: **A**. signaling protocol used in the differentiation process. After two days of culture, cells are in an epiblast-like state. At that point morphogens are modified to induce differentiation. The different signaling regimes correspond to temporal perturbations of three morphogens: CHIRON, FGF and PD. Data collection times are specified on the top row. **B**. The landscape that affects the cells evolution in time changes according to the combination of signals present in the media at each time point. **C**. Cell-state proportions computed from FACS data (Experiment, left) compared to average performance of the fitted model (Simulations, right) at 7 time points (D2-D5) for 6 different signals (No C, *…*, C2-5). These show the effect of the morphogens on cell identity distributions and the performance of the model reproducing them. **D**. PCA projection of the vectors of proportions for all time points and states considered (35-dimensional). Dots correspond to simulated proportions obtained from the model, filled shapes correspond to different replicates of the same experiment. Colors correspond to different signaling regimes as indicated.

We have used five different types of classifiers: Linear Discriminant Analysis (LDA), Support Vector Machines (SVM), K-Nearest Neighbors (KNN), spanning forests and Convolutional Neural Networks (CNN) with hyperparameter optimization performed in MATLAB®. We analysed the different classifiers with data sets of different sizes and decided to illustrate the results using an SVM classifier taking 1000 repetitions of each signal considered both for training and test (see SI, section 1B for details). The entries of a classifier’s confusion matrix are the proportions *p*_*ij*_ of responses produced under signal *S*_*i*_ that are classified as responses arising under signal *S*_*j*_. From this confusion matrix the MI between *S* and *Ŝ* is given by

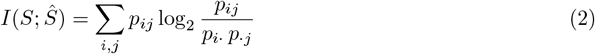

where 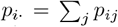 and 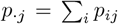 (see [4]). For the example presented in Fig. 1 the confusion matrix (normalised so the rows add up to one) for a classifier trained to separate the four signaling regimes and the corresponding contribution of each term to the sum in (2) are given in (3) and (4), respectively.

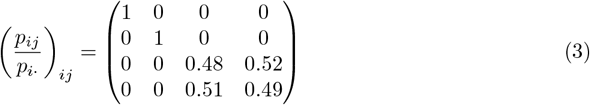

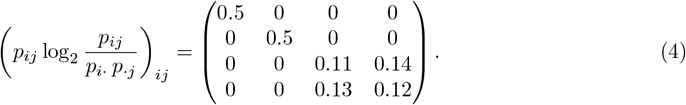

Hence, Pot = 2^1.5^ = 2.83.

## 4 Potency of the neural-mesodermal differentiation system

We demonstrate our approach using data from the in-vitro differentiation system of epiblast-like cells into neural and mesodermal progenitors considered in [11, 23, 25, 1] and modelled in [15]. In the experimental set-up, mouse stem cells that have been cultured in-vitro and differentiated into an epiblast-like state are considered. Afterwards, these cells are cultured under different signaling regimes defined by a temporal combination of the small molecule agonist of the WNT signaling pathway CHIR99021 (CHIRON), FGF2 (FGF) and the FGF inhibitor PD173074 (PD) as detailed in Fig. 2A. These morphogens induce the epiblast-like cells to differentiate into anterior neural, posterior neural or paraxial-mesodermal progenitors, going through specific transitions including a caudal epiblast identity. The differentiation process is selectively stopped at several time points along the differentiation pathway and single-cell FACS analysis is performed on the samples. Hence, the resulting experimental data consists of a 5-dimensional vector of protein expression for the 5 marker genes in each cell at each time point and for each signaling regime.

The state of each cell is characterised using the protein expression level of 5 well-studied marker genes: Nkx2.2, Olig2, Pax6, Nkx6.1, Sox2. The cells are clustered in the multidimensional protein expression space by fitting a Gaussian Mixture Model (GMM) to this data (see [15]). The fitted GMM gives clusters that correspond to either one of the cell states or a transitioning state between them. This clustering is then used to assign an identity to each single cell. For each time point and each signaling regime, we can then compute the proportion of cells in each state (Fig. 2C, Experiment). These proportions quantitatively characterise the outcome of each experiment and they are used to fit a mathematical model (Fig. 2B) that is able to reproduce the cell-type proportions at the different sample times under the different temporal signaling profiles (Fig. 2C and SI).

The mathematical model we consider for this system was presented in [15]. It is a landscape based model (Fig. 2B) that consists of 5 attractors, one for each differentiated cell state in the biological system. All trajectories start at the Epiblast attractor and follow the trajectories determined by the geometry of the landscape that changes according to the signaling regime that is being simulated (Fig. 2B).

The fitting of the model was performed using an ABC SMC method that provides a distribution for the values of the realistic parameters [15, 3]. This together with the stochastic nature of the simulations results in stochastic variability in the simulation results that can be compared to the experimental variation encountered when repeating the same experiment at different times or when they are performed by different laboratories (Fig. 2D). We use the simulations obtained from this model as a substitute for the generation of experimental repeats of the system. As mentioned above, a high number of repeats of the decision process are needed to compute the potency of the system, which would be infeasible to obtain only from experiments.

In this setting, the generic signal that we described in Section 1 is materialised as the temporal signaling profile applied to each sample and the response is given by a vector containing the proportion of each cell type for each time point. We consider 5 different cell types that correspond to the five attractors in the landscape model obtained in [15]. The model is stochastic and for each simulation we assign an identity to the simulated cells based on a GMM model fitted to the ensemble of simulated data (see [15] for details). Since we are considering 5 cell types and 7 time points, the response to each experiment is a 35-dimensional vector.

We begin by considering the set of 6 signals corresponding to the different durations of CHIRON addition from [15]. Figure 2D shows the projection of the corresponding proportion vectors into the first two principal components of the data. We observe the separation of the distributions of the different signaling regimes in the PC space. Using the method described in Section 3, the computation yields a potency approximately equal to 4.76 (see SI for details). This tells us that 4 signals can be distinguished, with some significant extra remaining information to distinguish between other signals. We call the difference between the number of signals and the potency the *potency gap* which in this case is 1.24. Such a gap is caused by *confusion* whose quantification we discuss below.

Including all the basic morphogen signals of [15] (i.e. including those that change FGF and/or PD levels) gives 11 signals. With this full set of 11 signals this system has a high potency of 9.533, and from the confusion matrix (Fig. 3B), it is clear that the reduction from 11 is due to the confusion introduced by the signaling regimes CHIRON days 2 to 4 (C2-4) and CHIRON days 2 to 5 (C2-5). If C2-4 is taken out, the potency rises to 9.843, which is remarkably close to the maximum of 10. The confusion matrix clearly shows that the classifier has no difficulty identifying 9 of the signal responses but gets the response to C2-4 wrong nearly 60% of the time and to C2-5 right only 61% of the time (Fig. 3B).

**Figure 3:**
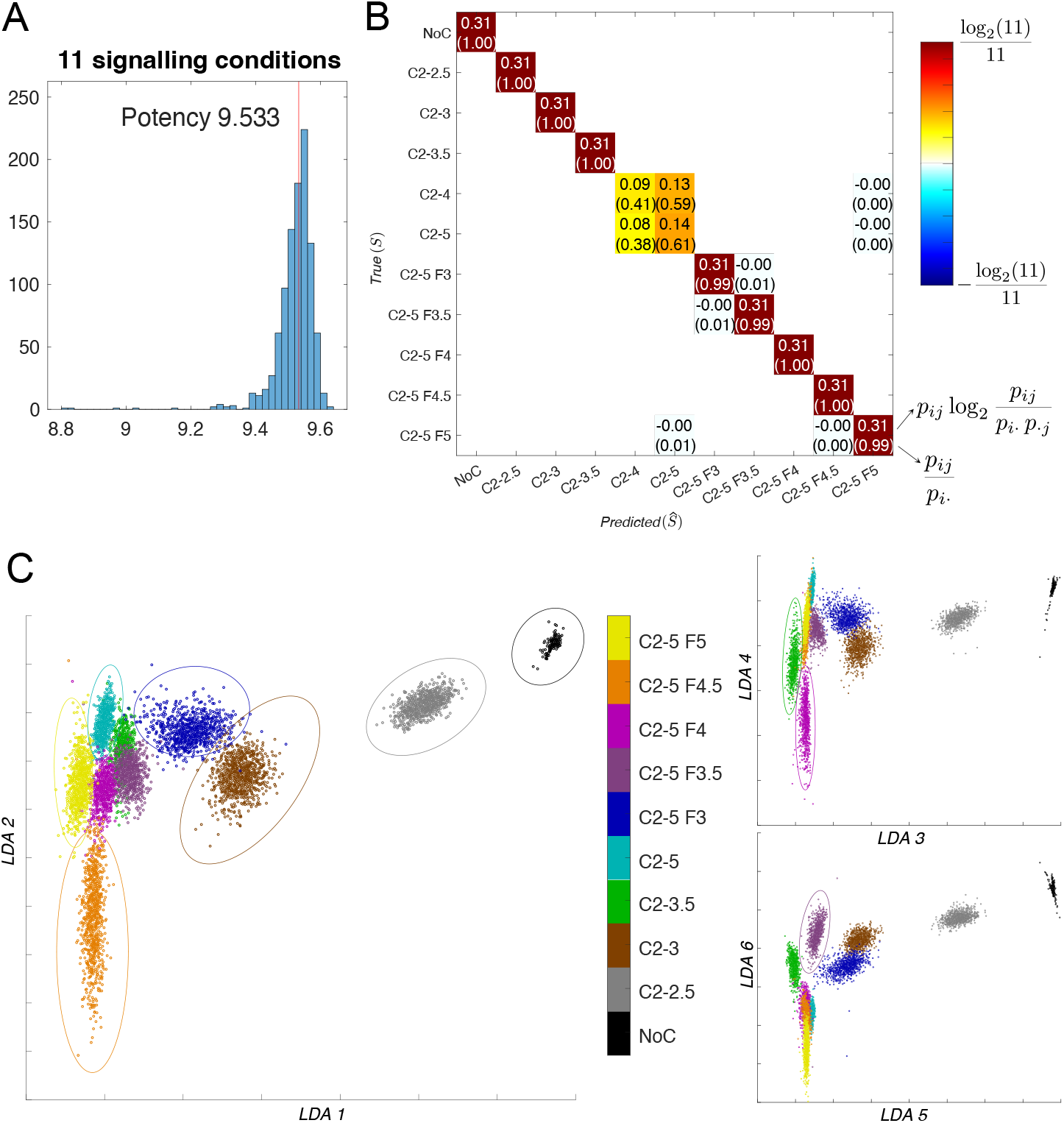
Potency computations for the neural-mesodermal differentiation system. **A**. Distributions for the 1000 repeats performed for the computation of the potency for all 11 signals considered. The potency is the median value marked with a red vertical line.**B**. Confusion matrix for the basic 11 conditions in [15] obtained from a classifier performing close to the potency value. The confusion between C2-4 and C2-5 conditions is apparent. The matrix shows the terms in (2) together with normalised values in the confusion matrix in brackets (normalised by rows). Rows correspond to true signals and columns correspond to predictions by the classifier. The color code is derived from the contribution to information as shown in the color bar. **C**. LDA projections of the 10 experimental conditions specified. The chosen projections show the separation of the 10 sets of responses.

**Figure 4:**
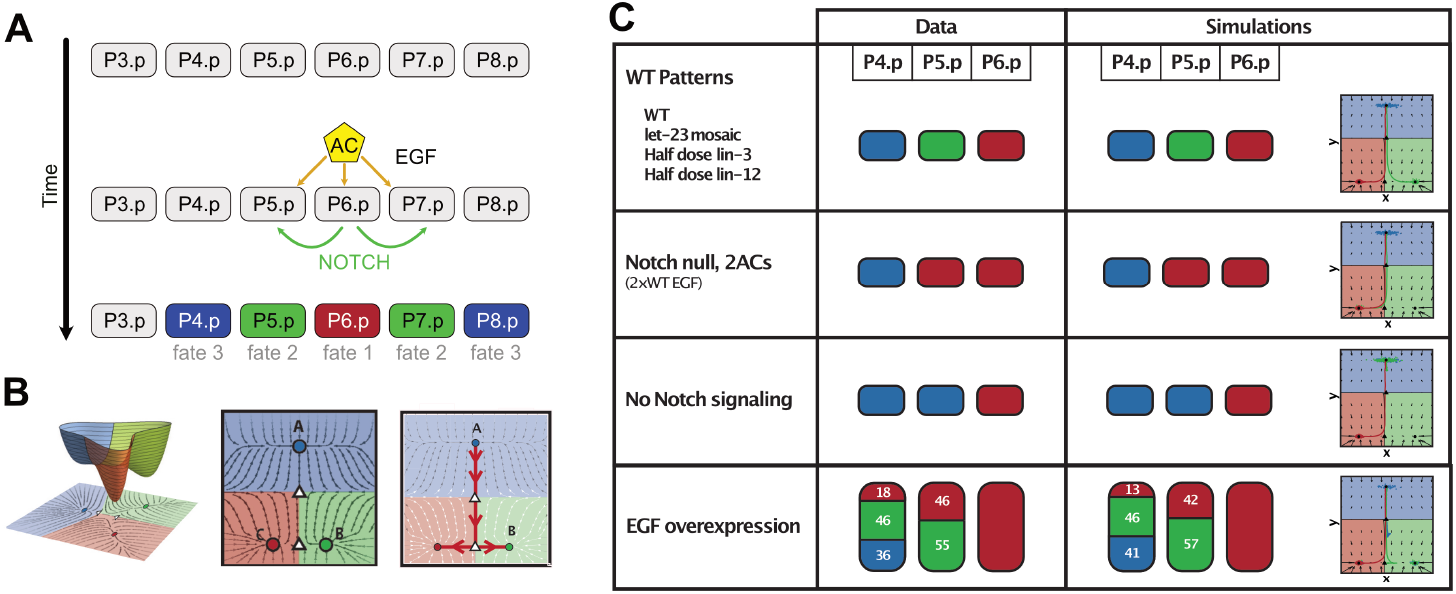
*Adapted from [3]*. **A**. Anchor cell (AC) induces vulval precursor cells (VPCs, P3–8.p) to differentiate into three different cell types: primary fate (red), secondary fate (green) tertiary fate (blue). The pattern is controlled by two signals, EGF from the AC (red arrows) and paracrine Notch (green arrows). VPCs are colored according to the WT pattern. P3.p is not colored because it is not included in our study. **B**. Model chosen to fit the data: 2-dimensional landscape with two decision points: vulval vs non-vulval fate (top saddle) and primary vs secondary fate (bottom saddle). This is the binary flip with cusp landscape. **C**. Model performance results. First column lists the conditions considered to train the model, second column shows the data on experimental proportions used and third column shows the quantitative results of the simulations of the fitted model together with an illustration of the model in each case.

We summarise the different results obtained in table 1 where we show how many signals can be identified in the different cases considered. Examples of the confusion matrices obtained are shown in the SI section 2A. Details for the list of conditions in each case are given in the caption of Table 1.

**Table 1:** The potency as a function of the number of signaling regimes. The initial 5 signaling regimes we consider are NoC, C2-2.5, C2-3, C2-3.5, C2-5 (see Figure 2). For 6, we consider the previous 5 and C2-4 and for 10 signals we consider all signaling regimes except C2-4. For 11 we take all the experimental conditions. See SI, section 11.4 for details.

| Number of signaling regimes | 5 | 6 | 10 | 11 |
| --- | --- | --- | --- | --- |
| Potency | 5 | 4.76 | 9.843 | 9.53 |
| Standard deviation | 0.0011 | 0.0014 | 0.0413 | 0.0635 |

Figure 1, through a simple example, introduces the concept of confusion — that adding new signals can reduce potency when the response to the new signal overlaps with the responses to existing ones; and this overlap diminishes the system’s overall discriminatory power, despite the presence of an additional signal. To quantify and better understand the confusion and the potency gap in this more complex context we compare the confusion matrices to the *D*_*KL*_ divergences of the distributions underlying the clusters of the 35-dimensional data points that define the responses. If we look at the responses in terms of the 35-dimensional vector associated to each response then, for each signal *S*, the responses form clusters that can be well represented as a sample from a roughly MVN distribution in a 35-dimensional space.

Moreover, if we can find a 2-dimensional plane in this 35-dimensional space of the response to the signals in which the projection of one of these clusters is disjoint from all the other clusters, then we can deduce that the cluster is disjoint for the whole domain. Thus by looking at projections of the responses to various linear planes, derived using the LDA method, we see in Fig. 3C that the 7 signals NoC, C2-2.5, C2-3, C2-5, C2-5 F3, C2-5 F4.5 and C2-5 F5 are effectively non-overlapping (Fig. 3C(left)), that, similarly, C2-3.5 and C2-5 F4 are disjoint from the rest in the projections shown in Fig. 3C(upper right) and, finally, that C2-5 F3.5 is disjoint from the rest in Figure 3C(lower right). On the other hand, the distributions of the responses to signals C2-4, C2-5 are overlapping to each other (see Figure 2D), which leads to the confusion recorded in the matrix presented in Figure 3B. It follows that 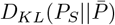 is close to log_2_ 11 if *S* ≠ C2-4, C2-5 and 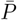 is the equal weight mixture of the *P*_*S*_. Consequently, the confusion causes a drop in *J* of 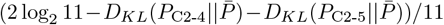. In the extreme case where *P*_C2-4_ = *P*_C2-5_, this is 2*/*11 because then 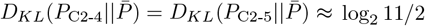 since *P*_C2-4_ and *P*_C2-5_ are approximately disjoint from the other *P*_*S*_.

## 5 Potency of the worm vulva patterning system

We now consider a self-organising developmental system where the *signals* correspond to complex experimental perturbations that do not necessarily affect a single signaling molecule or pathway in isolation. In this context, the potency measures the number of perturbations that give rise to distinguishable outcomes, which is useful not only for experimental design (by identifying perturbations that provide new information), but also as a way of measuring the potential of the underlying GRN to produce other clearly distinguishable responses, as these perturbations can be interpreted as mimicking or replacing natural signals.

In the normal development of the worm C. elegans vulva, six vulval precursor cells (VPCs), labeled P3–8.p, develop into three different fates: primary (1), secondary (2) and tertiary (3). They are partially differentiated and situated in a row along the anterior-posterior axis on the ventral side of the larva. In the WT, there is a precise pattern of cells fates in which cells P4–8.p adopt the fates 3,2,1,2,3 with very high probability. Although there is an extensive amount of data available, the sequence of events that promote this precise patterning is still unknown. The fate of each cell depends on an EGF signal received from an anchor cell (AC), positioned in the gonad of the larva, and the paracrine Notch signaling between the VPCs. The cell pattern therefore depends on the distance of each VPC to the AC, the paracrine signaling between them, and the tightly coordinated interplay between signaling dynamics and the progression of cell differentiation.

Building on the pioneering work of [7, 6], [3] developed a gene-free landscape model fitted to a large amount of experimental data which enabled the calculation of a posterior distribution of the parameters of the model. Data was used from a total of 21 different experimental setups where perturbation involving excess and reduced EGF, reduced, ectopic and null Notch, AC ablation and perturbation of EGF Notch receptors and ligand gave various outcomes (Table 1 in [3]). These experiments were repeated multiple times and the results for each is expressed as the proportion of each of the three fates observed for each of the three cells P4-6.p (Table 1 in [3]). This provides a large amount of data on the summary statistics of this large set of experiments. A landscape model was developed that could be used to simulate the data under the different experimental perturbations. This was fitted to the data using the ABC SMC method and used to generate predictions for future experiments.

Here we take this estimated landscape and posterior distribution for the parameters and estimate its potency. We restrict ourselves to the 9 conditions described in Figure 5A (see caption) for this computation. We sample from the posterior distribution of the parameters, simulate the system using the model in [3] and assign fates to the cells according to the three attractors of the model. We disregard the spatial information of each cell when computing the response to each input signal. We summarise the different results obtained in Table 2 where we show how many signals can be identified in the different cases. For each number of conditions we made an initial exploration of the potency by testing all classifiers in all possible combinations of *n* different signals, for varying values of *n*. Using training data we determined the set of signaling regimes that could give the maximal outcome and then performed the potency computation for that set as described above (see SI, Section 11.4). We see that although the number of experiments can be as high as 9, under any criterion the potency is limited to around 4 or 5 suggesting significant redundancy in the experiments with respect to identifying further distinguishable signaling regimes. Moreover, we computed the capacity for each of these systems by optimising the probability *P* (*S*) of the different signals. That is, for potency computation we assume they are equally likely while for capacity computation we searched over different probabilities (weight assigned to each perturbation in the discrete distribution *P* (*S*)) to find the maximum MI, *I*_*max*_ = max_*P* (*S*)_ *I*(*S*; *Ŝ*), and the capacity 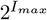 (see SI for details). We note that we get capacity values above 5 but the differences are fairly small.

**Table 2:** The potency and capacity as a function of the number of signaling regimes for the worm vulva patterning system.

| Num. of signals | 5 | 6 | 7 | 8 | 9 |
| --- | --- | --- | --- | --- | --- |
| Potency | 4.639 | 5.059 | 4.841 | 4.624 | 4.362 |
| Std. for Potency | 0.0164 | 0.0112 | 0.0108 | 0.0150 | 0.0128 |
| Capacity | 4.656 | 5.124 | 5.147 | 5.181 | 5.227 |
| Std. for Capacity | 0.0126 | 0.0130 | 0.0085 | 0.0104 | 0.0096 |

**Figure 5:**
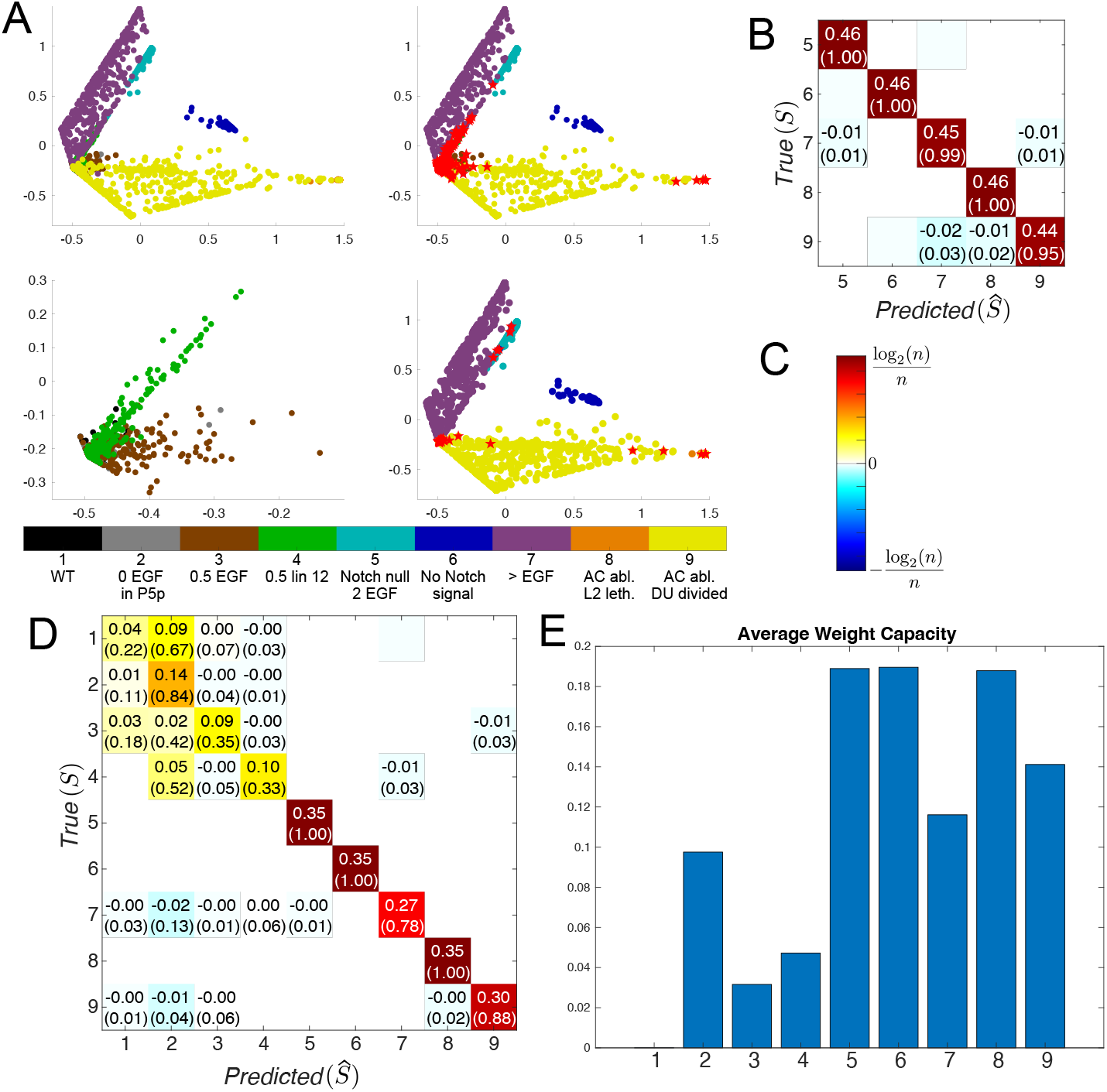
Potency and capacity computations for the vulval patterning system. **A. (Bottom panel)** The colours correspond to the different experiments which are: (1) Wild type, (2) let-23 mosaic (No EGF receptors in P5/7.p), (3) Half dose of lin-3 (Half EGF ligand), (4) Half dose of lin-12 (Half Notch receptor), (5) Notch null with double EGF (2 anchor cells), (6) No Notch signaling with WT EGF, (7) JU1100 (overexpression of EGF ligand), (8) L2 lethargus (anchor cell ablation), (9) Anchor cell ablation at DU divided. The scatter plots are PCA projections of the proportions of cells in the first two principal components. Points are coloured according to which experiment they arise from. **(Top left panel)** The 9 conditions considered from [3]. **(Top right panel)** Same conditions but includes red stars for the errors of the classifier for this set of conditions. **(Bottom left panel)** Shows the area where most confusion happens enlarged. **(Bottom right panel)** Shows the conditions that achieve the highest potency, as considered in panel B, together with the mistakes made by the classifier marked with a red star. **B**. Confusion matrix for the 5 conditions that achieve the highest potency obtained from a classifier performing close to the potency value. **C**. Color code for the matrices in the figure where *n* is the number of signals considered in each case. **D**. Confusion matrix for all 9 condition condidered obtained from a classifier performing close to the potency value. **E**. Average weights obtained for each condition in the performance of the 1000 repeats of the capacity computation.

For the vulval patterning system the confusion matrix in Fig. 5D makes it clear that there is much more confusion between responses. For the full set of 9 signals the first 4 confuse each other and 7 and 9 have significant interactions with one or two of the first 4 signals (Fig. 5A,D). Reducing the signals to 5, 6, 7, 8, 9 reduces the confusion to a level that can perhaps be ignored with classifier errors reduced to a few percent but adding a sixth signal can increase the potency to approximately 5.06. The maximum capacity which occurs with 9 signals at 5.22 is only a little larger.

These results suggest that this system is significantly more constrained than the Neural-Mesodermal differentiation system above in that it seems much harder to generate new perturbations that are distinguishable from the initial ones. We conjecture that this is because the system is more complex in that it involves the coupled dynamics of decision making by three cells that see different signals. The landscape of each of the three cells involves three attractors and two saddles and the trajectories by which the coupled system reaches the final state are complex.

## 6 Discussion

The paper introduces a new way of considering the flow of information in decision-making systems from signals to patterns. It seems reasonable to consider the GRNs that govern the sort of decision making underlying development as signal-dependent dynamical systems that provide the routes that guide cells from their precursor states through to their end fates. As such, to function and produce a diverging distribution of cell states they must have the ability to guide cells along diverging paths and to control the number and spatial location of the cells transitioning along such paths. From this viewpoint the ability of the signals to change the traffic flow and resulting end patterns is a primary characteristic of the system and it is this aspect that our potency attempts to quantify.

As discussed in [16] the modeling approach taken in [3, 15] is a natural way to understand development, and such models are effective at reproducing the decision making behaviour at the population level, reproducing experimental data and giving quantitative predictions of the behaviour of the considered system. The approach presented here needs a high number of replicates of the decision process both for training and testing, which is not feasible to obtain with the available experimental techniques. Landscape models easily provide the necessary replicates of the data and enable us to approximate the information transduction in the real experimental or in-vivo system. The accuracy of the potency approximation will always depend on the model’s accuracy in approximating the real TRP and its experimental variability.

It is worth emphasising at this point that we are not able to estimate the probability *P* (*S*) in-vivo, which is why we work with possible experimental distributions of the signals. Moreover, there is a question to be discussed about how this probability would be chosen by biology to guarantee the success in embryo development. Here we also estimate the maximal possible value of potency, but in a real organism some confusion, that is existence of non-distinguishable signals, may be necessary to guarantee the robustness of the output obtained. It is reasonable to consider desirable that small perturbations in the signal produce no difference in the outcome, so that the individual develops normally in spite of defects in the originating signals.

While the response patterns of our examples are relatively simple, it is clear that such a notion of potency can in principle be extended to a much richer set of responses that include both spatial and temporal patterns. One only needs to extract the appropriate features that summarise the responses to then use as data for the training and testing of the corresponding classifier.

As mentioned above, one needs a large number of measurements corresponding to several populations exposed to the same signals in order to obtain several proportion vectors corresponding to the same signal *S* so as to obtain a distribution of those, which explains the need for a realistic model in order to generate such data. Recent technologies (e.g. use of organoids) make the availability of replicates more common and could provide such experimental data in the near future.

This approach can also be useful in the context of experimental design and optimisation. As shown in our second example, where we considered experimental perturbations rather than genuine signals, the computation of potency in this context can be also seen as a quantification of what new experiments could lead to the most interesting and informative outcomes. The use of landscape models to predict new experimental outcomes was done in [7, 6, 10, 3] although no such optimisation was considered there.

## Acknowledgements

This research was supported in part by grant NSF PHY-2309135 to the Kavli Institute for Theoretical Physics (KITP), and UK Engineering and Physical Science Research Council (EPSRC) (grants EP/P019811/1 and EP/T031573/1). The authors thank James Briscoe and David Bruckner for useful discussions along the way.

## References

[1] R. Blassberg, H. Patel, T. Watson, M. Gouti, V. Metzis, M. J. Delás, and J. Briscoe. Sox2 levels regulate the chromatin occupancy of wnt mediators in epiblast progenitors responsible for vertebrate body formation. Nature cell biology, 24(5):633–644, 2022.

[2] Clive G Bowsher, Margaritis Voliotis, and Peter S Swain. The fidelity of dynamic signaling by noisy biomolecular networks. PLoS computational biology, 9(3):e1002965, 2013.

[3] E. Camacho-Aguilar, A. Warmflash, and D.A. Rand. Quantifying cell transitions in c. elegans with data-fitted landscape models. PLOS Computational Biology, 17(6):1–28, 06 2021.

[4] S.A. Cepeda-Humerez, J. Ruess, and G. Tkačik. Estimating information in time-varying signals. PLOS Computational Biology, 15(9):1–33, 09 2019.

[5] Raymond Cheong, Alex Rhee, Chiaochun Joanne Wang, Ilya Nemenman, and Andre Levchenko. Information transduction capacity of noisy biochemical signaling networks. Science, 334(6054):354–358, 2011.

[6] F. Corson and E.D. Siggia. Geometry, epistasis, and developmental patterning. Proc Natl Acad Sci U S A, 109(15):5568–5575, 03 2012.

[7] F. Corson and E.D. Siggia. Gene-free methodology for cell fate dynamics during development. eLife, 6:e30743, ec 2017.

[8] Thomas M Cover and Joy A Thomas. Information theory and statistics. Wiley New York, 2 edition, 2006.

[9] Julien O Dubuis, Gašper Tkačik, Eric F Wieschaus, Thomas Gregor, and William Bialek. Positional information, in bits. Proceedings of the National Academy of Sciences, 110(41):16301– 16308, 2013.

[10] Marine Fontaine, M. Joaquina Delas, Meritxell Saez, Rory J. Maizels, Elizabeth Finnie, James Briscoe, and David A. Rand. Dynamic landscape analysis of cell fate decisions: Predictive models of neural development from single-cell data. bioRxiv, 2025.

[11] M. Gouti, A. Tsakiridis, F. J. Wymeersch, Y. Huang, J. Kleinjung, V. Wilson, and J. Briscoe. In vitro generation of neuromesodermal progenitors reveals distinct roles for wnt signalling in the specification of spinal cord and paraxial mesoderm identity. PLOS Biology, 12(8):1–15, 08 2014.

[12] Adityanand Guntuboyina. Lower bounds for the minimax risk using f-divergences, and applications. IEEE Transactions on Information Theory, 57(4):2386–2399, 2011.

[13] Lauren McGough, Helena Casademunt, Milo š Nikolić, Zoe Aridor, Mariela D. Petkova, Thomas Gregor, and William Bialek. Finding the last bits of positional information. PRX Life, 2:013016, Mar 2024.

[14] D.A. Rand, A. Raju, M. Sáez, and E.D. Siggia. Geometry of gene regulatory dynamics. Proc Natl Acad Sci U S A, 118(38), 09 2021.

[15] M. Sáez, R. Blassberg, E. Camacho-Aguilar, E.D. Siggia, D. A Rand, and J. Briscoe. Statistically derived geometrical landscapes capture principles of decision-making dynamics during cell fate transitions. Cell Systems, 13(1):12–28, 2022.

[16] M. Sáez, D. A Rand, and J. Briscoe. Dynamical landscapes of cell fate decisions. Interface, 12(4), 06 2022.

[17] Jangir Selimkhanov, Brooks Taylor, Jason Yao, Anna Pilko, John Albeck, Alexander Hoffmann, Lev Tsimring, and Roy Wollman. Accurate information transmission through dynamic biochemical signaling networks. Science, 346(6215):1370–1373, 2014.

[18] Harish Shankaran, Danielle L Ippolito, William B Chrisler, Haluk Resat, Nikki Bollinger, Lee K Opresko, and H Steven Wiley. Rapid and sustained nuclear–cytoplasmic erk oscillations induced by epidermal growth factor. Molecular systems biology, 5(1):332, 2009.

[19] Gašper Tkačik and William Bialek. Information processing in living systems. Annual Review of Condensed Matter Physics, 7(1):89–117, 2016.

[20] Gašper Tkačik, Curtis G Callan Jr, and William Bialek. Information flow and optimization in transcriptional regulation. Proceedings of the National Academy of Sciences, 105(34):12265– 12270, 2008.

[21] Gašper Tkačik, Julien O Dubuis, Mariela D Petkova, and Thomas Gregor. Positional information, positional error, and readout precision in morphogenesis: a mathematical framework. Genetics, 199(1):39–59, 2015.

[22] Gašper Tkačik and Thomas Gregor. The many bits of positional information. Development, 148(2):dev176065, 2021.

[23] A. Tsakiridis, Y. Huang, G. Blin, S. Skylaki, F. Wymeersch, R. Osorno, C. Economou, E. Karagianni, S. Zhao, S. Lowell, and V. Wilson. Distinct wnt-driven primitive streak-like populations reflect in vivo lineage precursors. Development, 141(6):1209–1221, 2014.

[24] Margaritis Voliotis, Rebecca M Perrett, Chris McWilliams, Craig A McArdle, and Clive G Bowsher. Information transfer by leaky, heterogeneous, protein kinase signaling systems. Proceedings of the National Academy of Sciences, 111(3):E326–E333, 2014.

[25] F.J. Wymeersch, V. Wilson, and A. Tsakiridis. Understanding axial progenitor biology in vivo and in vitro. Development, 148(4), 2021.

[26] Yuhong Yang and Andrew Barron. Information-theoretic determination of minimax rates of convergence. Annals of Statistics, pages 1564–1599, 1999.

