## Supplemental Information for "Measuring developmental information encoded by a dynamical landscape"

March 4, 2026

#### 1 Methodology

In this section we provide the general methodology used to produce results in this manuscript.

##### 1.1 Estimating potency for a set of signals

First we provide the general algorithm used to produce estimates of mutual information and potency for a given set of signals. Here we assume that the data used for training and testing are labelled by a given set of signals.

**Algorithm 1:** Estimating potency for a signals set.

```
Input: Class of classifiers 𝒳, training data, testing data, L positive integer
1 for ℓ = 1, 2, ..., L do
2     Train a classifier χ ∈ 𝒳 using the training data;
3     Use χ to predict the labels of each testing sample;
4     Use the predictions to compute the confusion matrix (p_ij);
5     Use this confusion matrix to compute I(S; Ŝ) as in Eq. (2) and store this as Î_ℓ;
6 end
Output: The sample median Î_med of Î_ℓ, ℓ = 1, 2, ..., L, and the estimated potency
        Pot^ = 2^(Î_med)
```

##### 1.2 Class of Classifiers and training

Here we provide details on the input class of classifiers $\mathcal{X}$ and the training of classifier, $\chi \in \mathcal{X}$, in step 3 of algorithm 1. For this we used MATLAB®2023a and its automatic hyperparameter optimisation option with objective function the cross-validation missclassification error. The classes of classifiers, $\mathcal{X}$, used are:

- Regularized linear discriminant analysis (LDA) using the MATLAB® function *fitcdiscr* with equal class/signal prior probabilities and the optimisation algorithm finding optimal regularisation parameters for the covariance matrix and the threshold of non-zero coefficient of response variables.

- Ensemble of binary support vector machine (SVM) classifiers with linear kernel using function *fitcecoc*. The optimisation considers different coding designs (one-versus-one and one-versus-all), the kernel scale and Box Constraint parameters.

- $k$-nearest neighbour (KNN) classifier using function *fitcknn* with the optimisation considering different distance metrics along with the $k$ parameter.

- Decision tree (Sp Trees) using function *fitctree* with the optimisation finding optimal values of the Minimum Leaf size.

- Neural Network using function *fitcnet* with the optimisation algorithm selecting between different activation functions, along with the number and size of hidden layers.

#### 1.3 Computation of the training and testing data

The training data are produced by drawing $K$ independent and identically distributed samples from the response distribution $P_S$, of each signal, $S \in \mathcal{S}$, with $\mathcal{S}$ a finite set of $n$ signals. The same process is used to produce the testing data. We give details of the probability distribution $P_S$ for our two example systems in section 2.1, 2.2.

In our investigations, we vary the sample size $K$ and the set of signals $\mathcal{S}$ and analyse the effect of these changes. See section 1.4 for the method used for the latter study. We also investigated whether computing the confusion matrix using predictions made for the training examples lead to different results than the algorithm described above, which uses independent testing data. The results for the Neural-mesodermal differentiation system showed overestimation of potency when testing data are not used, which is persistent for increasing values of $K$ (see Figures 3-7). This is not surprising given that training is optimising the classifier parameters using the training data. All the presented results, except from the above figures, used predictions made for the testing data as in Algorithm 1.

#### 1.4 Varying the set of signals

For a set of $N$ signals $\mathcal{S}$, we estimated maximum potency over the subsets of $\mathcal{S}$ of a fixed number, $n \leq N$. Below we provide the algorithm used for this investigation.

**Algorithm 2:** Estimating potency for fixed number of signals

**Input:**

Classes of classifiers $\mathcal{X}_1, \mathcal{X}_2, \ldots, \mathcal{X}_c$,;

training and testing data for set of signals $\mathcal{S}$ with $|\mathcal{S}| = N$,;

$n$ positive integer $\leq N$, $L$ positive integer

List all subsets, $\mathcal{S}_i \subset \mathcal{S}$, $i = 1, 2, \ldots, k$ of $n$ signals ($k = \binom{N}{n}$);

**for** $i = 1, 2, \ldots, k$ **do**

  Store the training and testing data with labels the signals in $\mathcal{S}_i$ as $R_{tr,i}$, $R_{te,i}$.

  **for** $j = 1, 2, \ldots, c$ **do**

    Apply Algorithm 1 with Input: $\mathcal{X}_j$, $R_{tr,i}$, $R_{te,i}$, $L$ positive integer;

    Store Outputs as $\mathrm{Pot}_{ij}$

  **end**

**end**

**Output:** $\max_i \mathrm{Pot}_{ij}$, $\mathrm{argmax}_i \mathrm{Pot}_{ij}$

##### 1.5 Computation of Capacity

The capacity for the channel $S \to \hat{S}$ is defined as the supremum of the mutual information $I(S; \hat{S})$ with respect to the probability function, $\mathbb{P}(S)$, of the signal $S$, i.e. $\sup_{\mathbb{P}(S)} I(S; \hat{S})$. Assumming that the signal is discrete and takes values $S_0, \ldots, S_n$ with probabilities $w_1, \ldots, w_n$, the channel capacity is

$$\sup_{(w_0,\ldots,w_n)\in\mathcal{W}^n} \sum_{i,j} w_i p_{j|i} \log_2 \frac{p_{j|i}}{\sum_{i'} w_{i'} p_{j|i'}}. \tag{1}$$

Here $\mathcal{W}^n$ is the $n$-Simplex, i.e. $\mathcal{W}^n = \{(w_1, \ldots, w_n) \mid w_i \geq 0, i = 1, \ldots, n, \sum_{i=1}^n w_i = 1\}$, and $p_{j|i} := P(\hat{S} = S_j \mid S = S_i)$ are the conditional probabilities of predicting the signal $S_j$ given that the response was generated under signal $S_i$. These conditional probabilities can be estimated by the normalised entries of the confusion matrix, with the normalisation making each row sum to 1.

The calculation of the above supremum can be solved numerically by a constrained optimisation algorithm. Here we derive this by the MATLAB® function *fmincon*. As in the computation of $I(S; \hat{S})$ we perform this optimisation a number of repetitions $L$ and take the median of the samples obtained as the approximation of the capacity. We store the weights in each repetition for analysis.

### 2 Results

#### 2.1 Neural-mesodermal differentiation system

The generation of simulated data for training and testing are described in Section 4 of the main paper.

We consider various scenarios. First we consider all available ($N = 11$) signals. Figure 1 provides an extended version of Figure 2(C), which is a comparison of simulated and experimental data of the response vectors for the $N = 11$ signals. Here we call signals the different conditions used for each experimental set up.

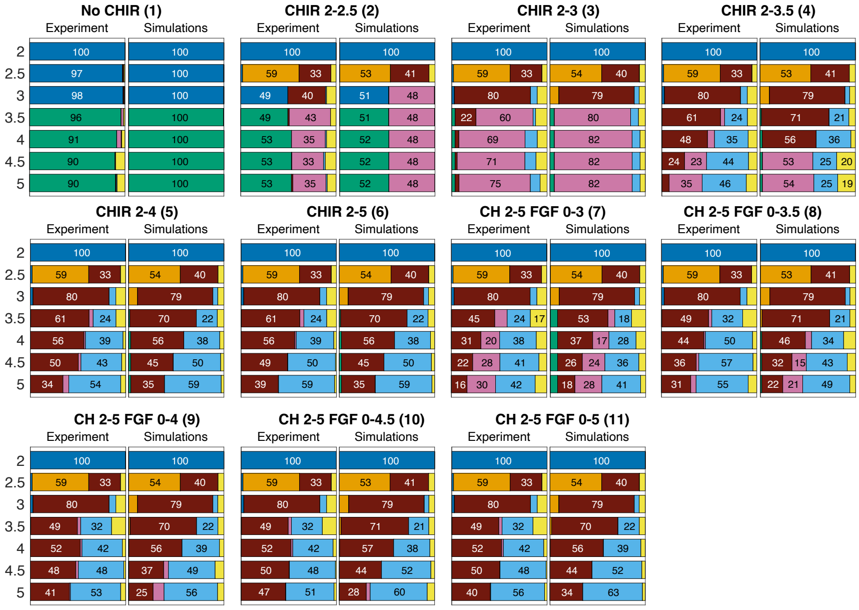


Figure 1: **Neuro-Mesodermal Progenitors data.** Mean proportions for all conditions from [15]. This is an extended version of Figure 2(C). See caption of Figure 2 for more details.

We draw $K = 500$ independent and identically distributed samples from the response distribution $P_S$, of each of those $N = 11$ signals. This is used as training data. We then repeat the same process to draw $K = 500$ samples for testing. We performed Algorithm 1 for each of the 5 classes of classifiers described in Section 1.2 with $L = 1$. The confusion matrices obtained are shown in Figure 2. We observe that all the classifiers struggle in differentiating between conditions 5 and 6. That was to be expected since these two conditions give virtually the same response as discussed in the main paper.

To simplify the remaining analysis, we remove condition 5 from the set of conditions, that is, we consider the remaining 10 conditions and run the algorithm using each of the classifiers above for the number of training samples $K \in \{100, 250, 500, 750, 1000, 2500, 5000\}$. We take $L = 1000$ for $K \in \{100, 250, 500, 750, 1000\}$. The distributions obtained are shown in the plots titled "Test" Figures 3 to 7.

We observe tighter distributions and increased (median) potency in as $K$ increases, but the differences are not substantial. Note also a substantial increase on average computational time for larger values of $K$. Comparing the different classes of classifiers, SVM gets the highest potency estimate in all cases, closely followed by the CNN classifier, with the Regularised LDA giving the lowest value.

After considering these results, we decided to perform the rest of potency computations using SVM classifiers with $L = 500$ and $K = 500$. That is, we apply Algorithm 2 using one class of classifiers, that is the SVM classifiers, $L = 500$, and training and testing data with $K = 500$ samples as explained in Section 1.3. For varying values of $n$, we derive the results presented in Table 1 and 2.

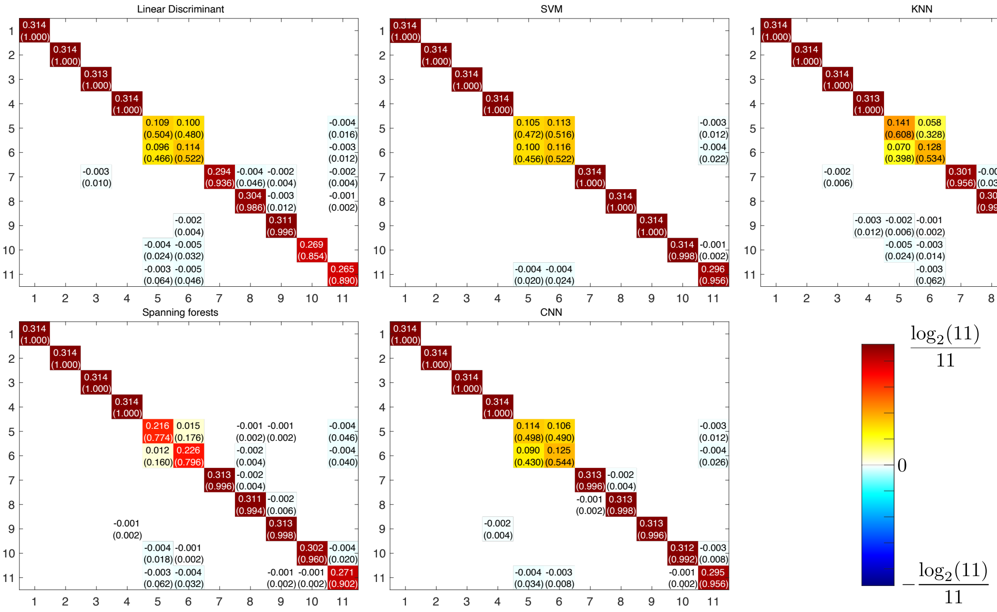


Figure 2: **Terms for computing the information in the neural-mesodermal differentiation system for the 5 classifiers considered.** All matrices in the figure show the terms in equation 2, together with normalised values in the confusion matrices in brackets (normalised by rows). Rows correspond to real signals and columns correspond to predictions by the classifier. The color code is derived from the contribution to information as shown in bottom right panel.

For the results in Figure 3(A-B) we applied Algorithm 1 for $L = 1000$.

#### 2.2 Capacity computation for the vulval system

Section 5 describes the procedures followed for generating data for the vulval system using the model in [3] and computing potency.

Here, we obtain the distribution of weights when repeating for $L = 1000$ times the optimisation performed for the computation of the capacity of the vulval system (see Table 2). The results are presented in Figure 8. We see that the conditions that maximise potency are those that were clearly differentiated from the others (shown by larger diagonal values in the confusion matrix) together with some of the ones that caused confusion (lower diagonal values in the confusion matrix). Notably, condition number 1 is almost always excluded since it is the most confused with the others.

We also consider the potency's dependence to the choice of different signals, when a fixed number $n = 8$ of signals is considered. We see a substantial variation between 3.71 and 4.9 is observed, with the highest potency achieved when signal 1 is dropped. The latter is in agreement with the results in Figure 8.

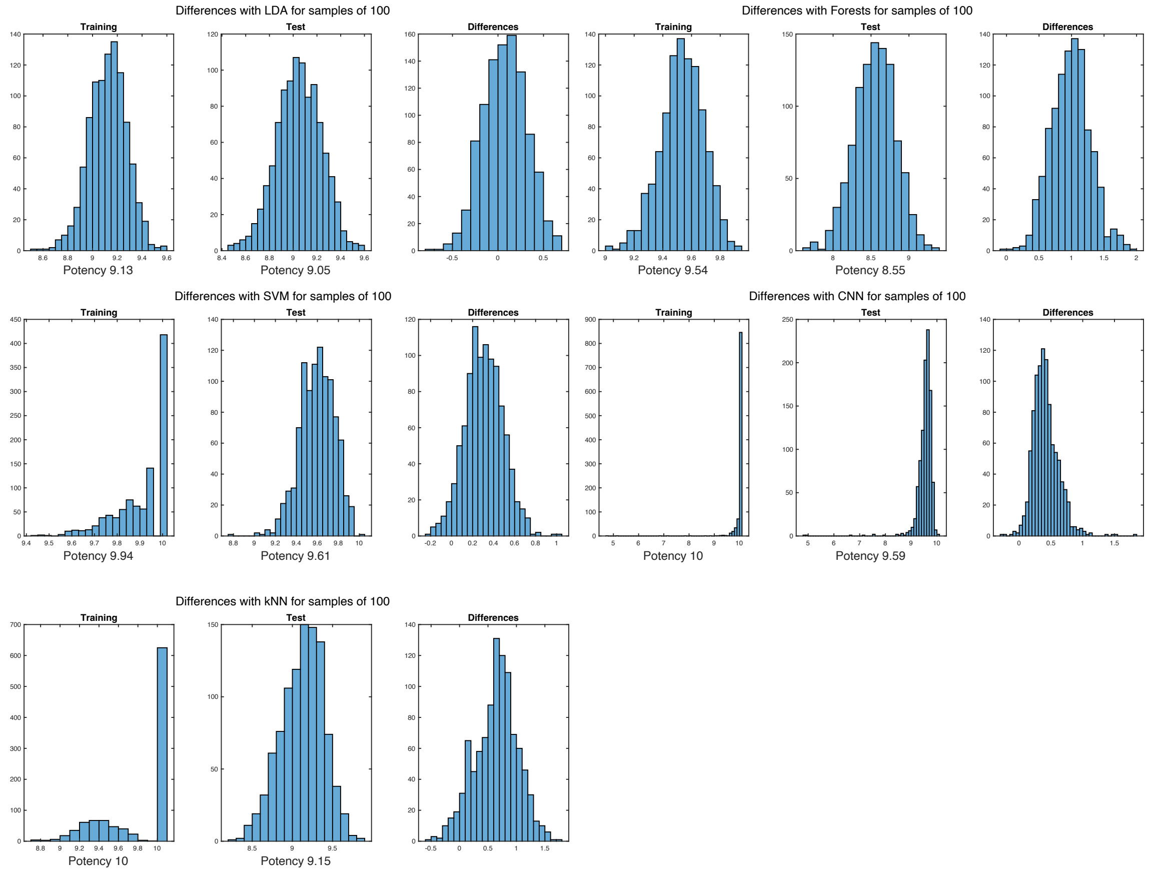


Figure 3: **Samples of 100** Potency computation results with $K = 100$. Each panel corresponds to a different classifier as specified in the title. For each panel, the rightmost subpanel corresponds to the computation using training data as test data, in the middle one the proper computation using test data disjoint from training data and the right one shows the difference between the two. Below the two first subpanels the median of the distribution corresponding to the potency value obtained is shown for comparison.

| Signals | Potency |
|---|---|
| 1 2 3 4 5 6 7 8 | 3.89 |
| 1 2 3 4 5 6 7 9 | 3.72 |
| 1 2 3 4 5 6 8 9 | 4.17 |
| 1 2 3 4 5 7 8 9 | 3.71 |
| 1 2 3 4 6 7 8 9 | 3.77 |
| 1 2 3 5 6 7 8 9 | 4.68 |
| 1 2 4 5 6 7 8 9 | 4.80 |
| 1 3 4 5 6 7 8 9 | 4.80 |
| 2 3 4 5 6 7 8 9 | 4.90 |

Table 1: Potency for different combinations of 8 signals of the vulval system. Only training data, and the SVM classifier are used here.

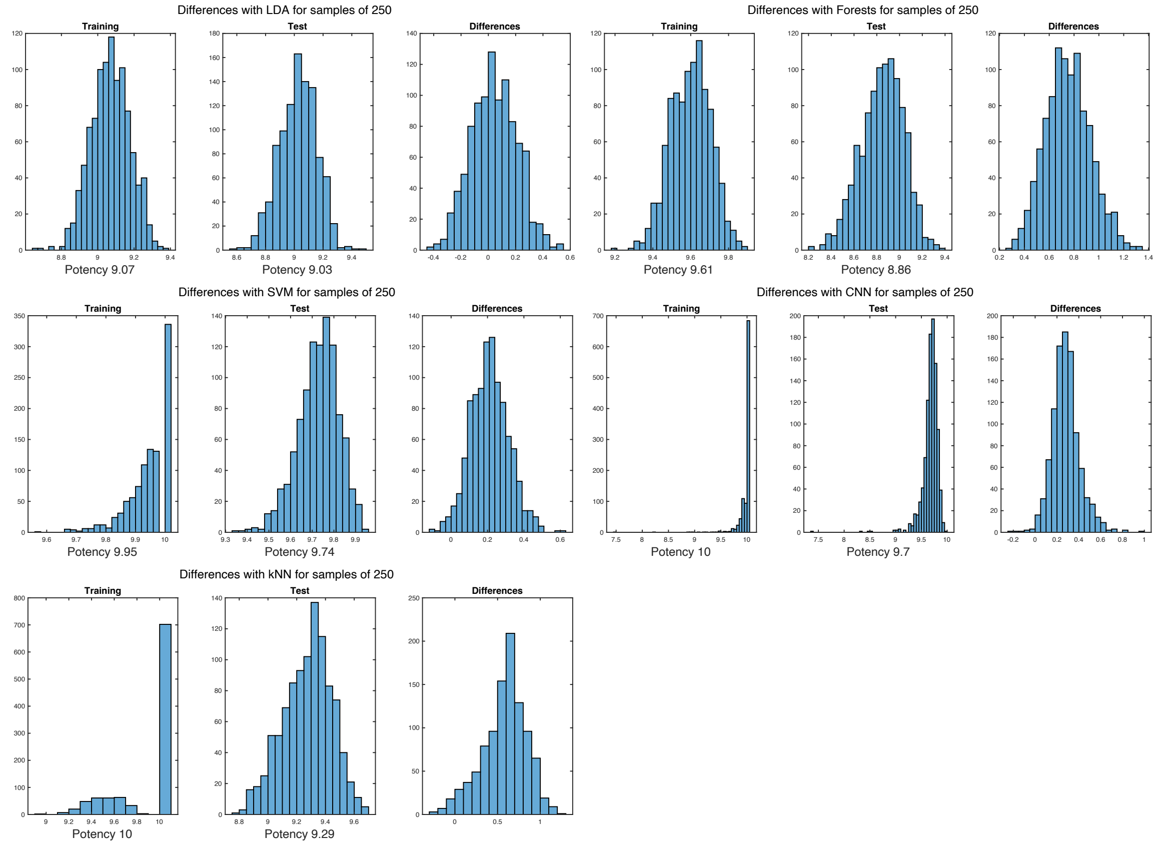


Figure 4: **Samples of 250** Potency computation results with $K = 250$. Each panel corresponds to a different classifier as specified in the title. For each panel, the rightmost subpanel corresponds to the computation using training data as test data, in the middle one the proper computation using test data disjoint from training data and the right one shows the difference between the two. Below the two first subpanels the median of the distribution corresponding to the potency value obtained is shown for comparison.

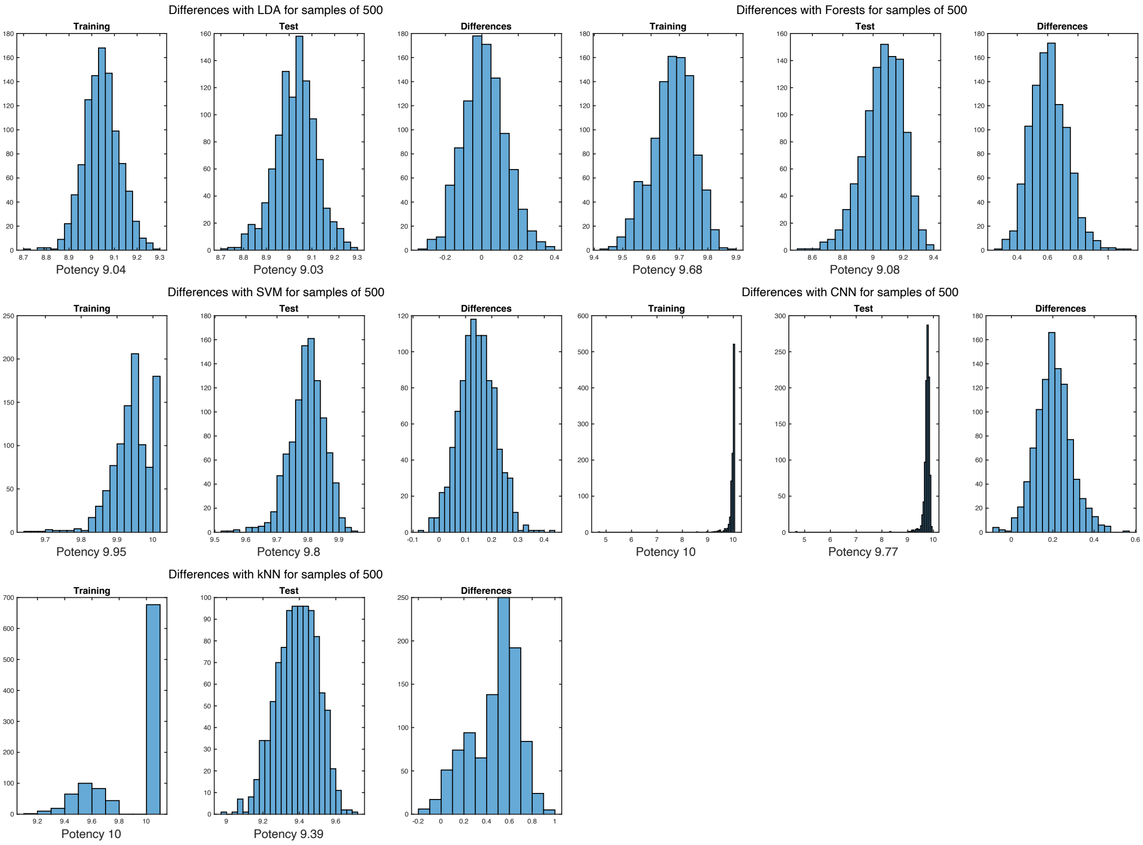


Figure 5: **Samples of 500** Potency computation results with $K = 500$. Each panel corresponds to a different classifier as specified in the title. For each panel, the rightmost subpanel corresponds to the computation using training data as test data, in the middle one the proper computation using test data disjoint from training data and the right one shows the difference between the two. Below the two first subpanels the median of the distribution corresponding to the potency value obtained is shown for comparison.

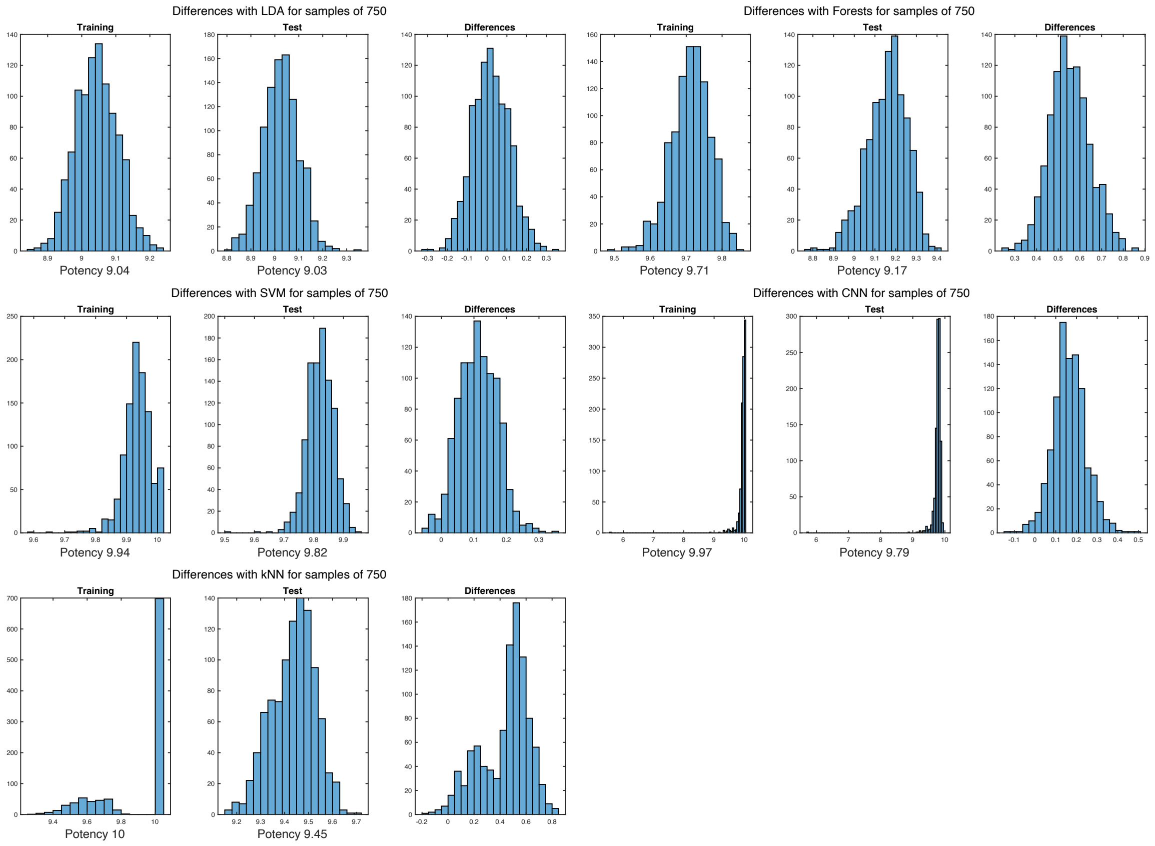


Figure 6: **Samples of 750** Potency computation results with $K = 750$. Each panel corresponds to a different classifier as specified in the title. For each panel, the rightmost subpanel corresponds to the computation using training data as test data, in the middle one the proper computation using test data disjoint from training data and the right one shows the difference between the two. Below the two first subpanels the median of the distribution corresponding to the potency value obtained is shown for comparison.

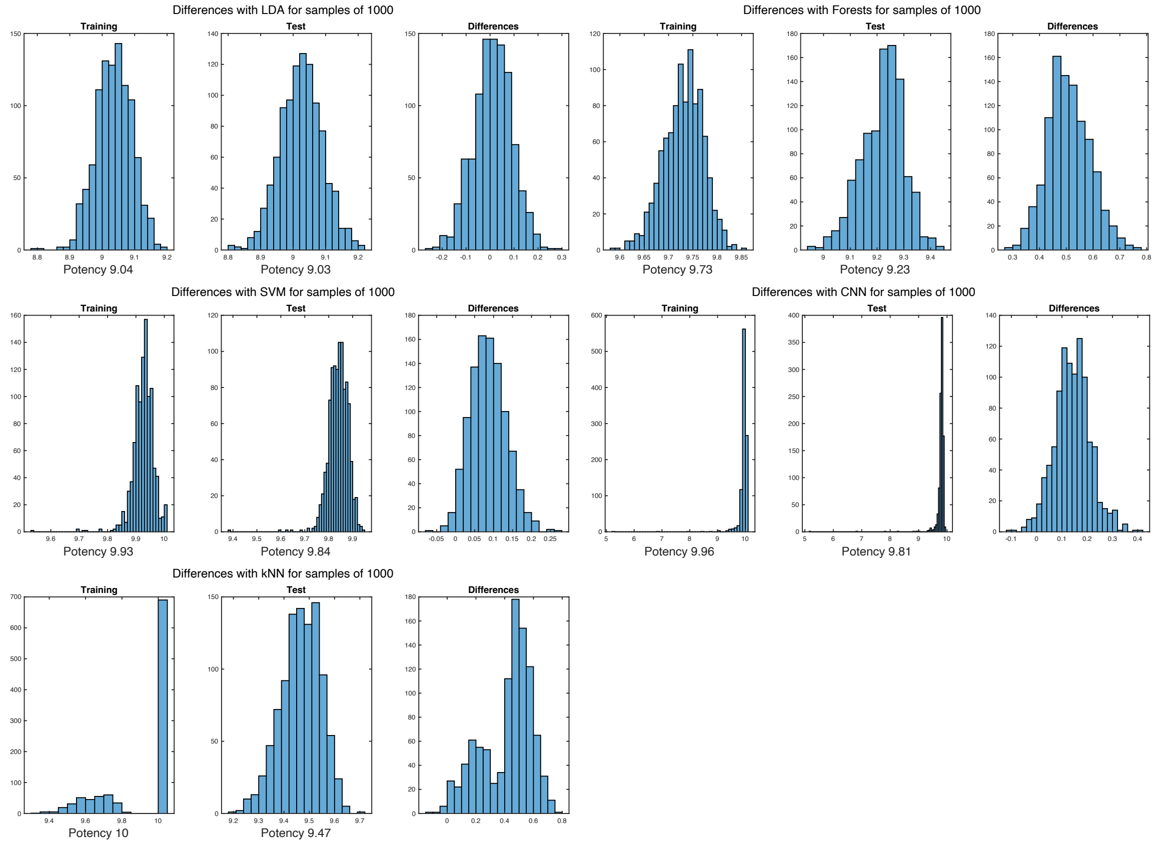


Figure 7: **Samples of 1000** Potency computation results with $K = 1000$. Each panel corresponds to a different classifier as specified in the title. For each panel, the rightmost subpanel corresponds to the computation using training data as test data, in the middle one the proper computation using test data disjoint from training data and the right one shows the difference between the two. Below the two first subpanels the median of the distribution corresponding to the potency value obtained is shown for comparison.

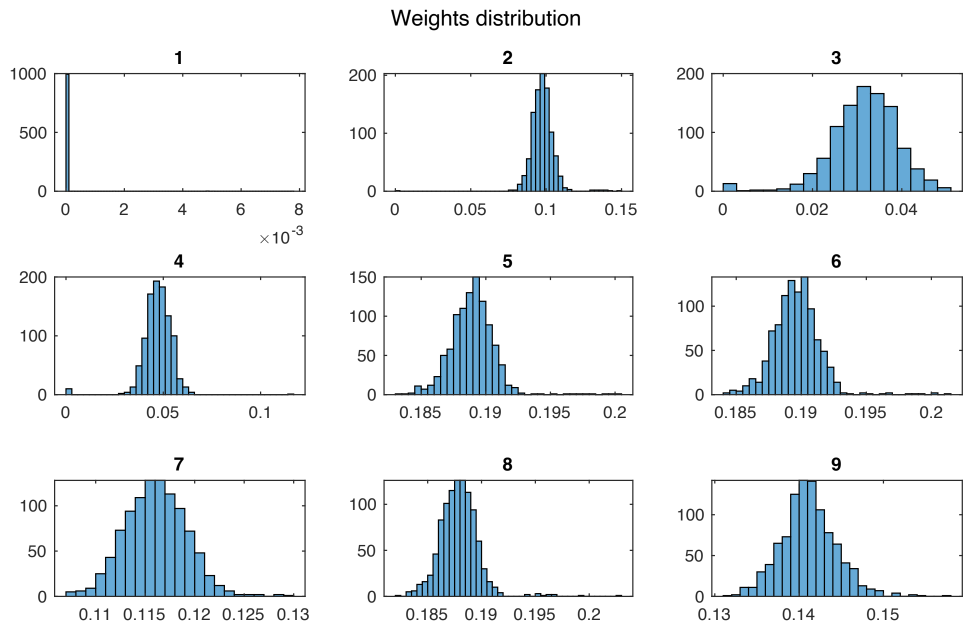


Figure 8: **Weights obtained in capacity computation.** The histogram of the weight of each of the 9 different signals considered in computing the capacity of the vulval system (see Table 2).

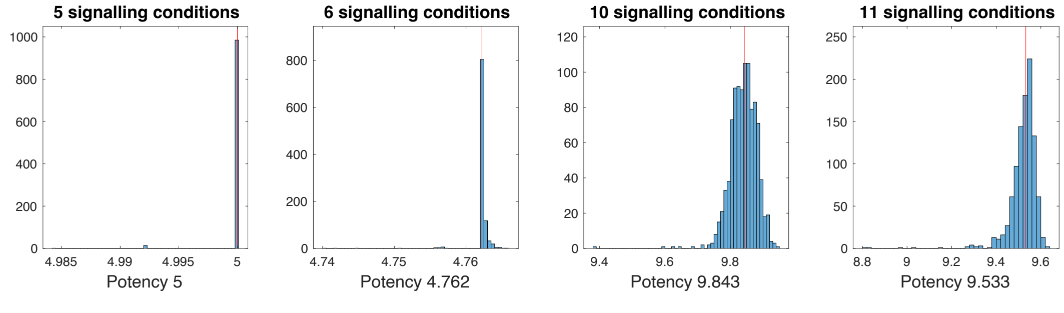


Figure 9: **Potency for the Neuro-Mesodermal Progenitors data** The computation was performed 1000 independent times. The median of the distribution is taken as the potency value as shown by the red line and the value is presented below. The first panel considers the 5 signals 1 to 4 and 6. The second panel considers the 6 signals 1 to 6. The third panel considers the 10 signals 1 to 4 and 6 to 11. The last panel considers all signals 1 to 11.

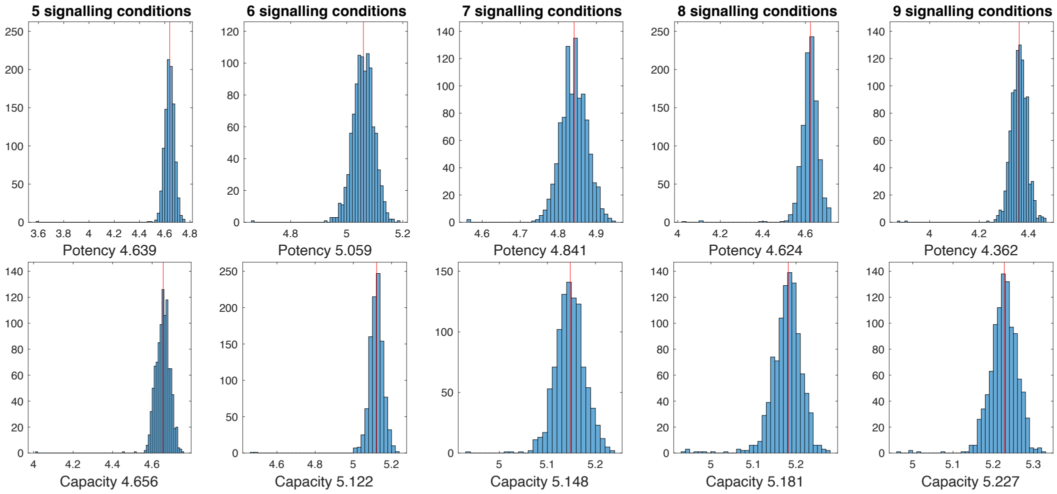


Figure 10: **Potency and capacity for the vulval progenitors data** The computation was performed 1000 independent times. The median of the distribution is taken as the potency (resp. capacity) value as shown by the red line and the value is presented below. The first set of two panels (vertically) considers the 5 signals 5 to 9. The second set considers signal 2 together with signals 6 to 9. The third set considers signal 2 together with signals 5 to 9. The fourth set considers signals 2 to 9. The last set considers all signals 1 to 9. These sets were chosen because they give the maximal values for that specific number of signalling conditions.
